# p190A/*ARHGAP35* and p190B/*ARHGAP5* proteins in endometrial cancer, a novel cancer-relevant paralog interplay

**DOI:** 10.1101/2025.10.16.682927

**Authors:** Mathilde Pinault, Capucine Heraud, Mélissa Correia De Oliveira, Véronique Neaud, Valérie Valesco, Valérie Prouzet-Mauleon, Béatrice Turcq, Jean-William Dupuy, Anne-Aurélie Raymond, Sabrina Croce, Frédéric Saltel, Valérie Lagree, Violaine Moreau

**Affiliations:** Univ. Bordeaux, INSERM, BRIC, U 1312, F-33000 Bordeaux, France; Department of Biopathology, Institut Bergonié, 33076 Bordeaux, France; CRISP’edit, TBMCore, Univ. Bordeaux, CNRS UAR 3427, Inserm US05, F-33000 Bordeaux, France; Plateforme Protéome, Univ. Bordeaux, F-33000 Bordeaux, France; OncoProt, TBMCore, Univ. Bordeaux, CNRS UAR 3427, Inserm US05, F-33000 Bordeaux, France

**Keywords:** Paralogs, RhoGAPs, endometrial cancer, synthetic lethality

## Abstract

Endometrial cancer is one of the main gynecological malignancies worldwide, with an estimated 320, 000 new cases annually. Several studies highlight *ARHGAP35* as a significantly mutated gene in these tumors. It encodes for the protein p190RhoGAP-A (p190A), which is a major regulator of the small GTPase family of proteins. *ARHGAP5* is a paralog of *ARHGAP35* that encodes the protein p190RhoGAP-B (p190B). By analyzing human endometrial cancer samples, we found a co-occurrence of mutations in *ARHGAP35* and *ARHGAP5* genes and we reported that both are less expressed at the mRNA level in tumoral samples compared to healthy tissues. We were interested in understanding the impact of p190A/B under-expression in endometrial cancer and the relationship between the two paralogs. To do so, we have used CRISPR/Cas9 technology to generate HEC-1-A knockout cells for p190A and p190B. We showed that removal of each paralog led to a similar actin remodeling phenotype with the formation of Cross-Linked Actin Networks (CLANs), dependent on the Rho/ROCK pathway. Moreover, proteomic analysis of p190A and p190B knockout cells highlighted similar affected cell functions. Finally, our study demonstrates a synthetic lethality between p190A and p190B where removal of both paralogs is deleterious in endometrial cancer cells, unveiling a potential actionable vulnerability.

## INTRODUCTION

Endometrial carcinoma is one of the most common malignancies of the female reproductive system and the 4^th^ most common cancer in the Western world (*1*). The vast majority (over 90%) of endometrial tumors are diagnosed in women over the age of 50. Historically, the Bokhman classification distinguished two main types of endometrial carcinomas: type I, considered low grade, estrogen-related, less aggressive and histologically composed essentially of endometrioid type, and type II that included the clinically aggressive carcinoma, non-related to hormones and histologically made of serous and clear cell type (*2*). The TCGA molecular classification divided endometrial carcinomas into 4 subtypes: the POLE-mut subtype, which is ultramutated and associated with POLE (encoding DNA polymerase Epsilon) mutations; the mismatch repair-deficient (MSI) subtype, which is hypermutated and associated with microsatellite instability; a subtype associated with TP53 mutations with high copy number variation (CNV); and finally a subtype with no particular abnormality and low CNV (*3, 4*). These four molecular classes are correlated with patient survival: while POLE-mutated tumors are associated with a good prognosis, TP53-mutated tumors have a poor one (*3*). With the development of new therapeutic strategies, survival rates for patients with endometrial carcinomas have improved considerably. However, many patients respond poorly to treatment and rapidly develop recurrence and metastasis. Therefore, understanding the underlying mechanisms of tumor cell proliferation, migration and invasion is important to explore new targets for treating patients with endometrial carcinomas.

Sequencing of tumor exomes has identified *ARHGAP35* as a gene significantly mutated in 14% of endometrial tumors (*5, 6*), placing *ARHGAP35* in the list of the 30 most frequently mutated genes in human cancers. The *ARHGAP35* gene encodes p190RhoGAP-A (p190A), which belongs to the family of GTPase-Activating Proteins (GAPs) that regulate the activity of small GTPases of the Rho family. Nevertheless, this gene appears in the list of strict paralogs, together with the *ARHGAP5* gene, which encodes p190RhoGAP-B (p190B) (*7*) (http://ohnologs.curie.fr/). Whereas *ARHGAP35* was recently studied in endometrial cancer (*8, 9*), the status and impact of its paralog remain unknown.

In most human cells, the two paralogs *ARHGAP35*/p190A and *ARHGAP5*/p190B are co-expressed (*10–12*). These proteins of 190kDa show over 51% sequence identity, and share the same domain organization with their GAP domain responsible for the GAP activity at their C-terminal. In addition, from the N-terminal to the C-terminal, they show a GBD domain for GTP Binding Domain capable of binding GTP, four FF domains that are involved in proteins interactions and a central domain composed of two pseudo-GTPase domains. Recently, we identified the PLS domain in p190A implicated in the auto-inhibition of the GAP activity and in the regulation of the protein localization (*13–15*). By regulating RhoA, p190 proteins are key regulators of the actin cytoskeleton organization. Both proteins co-localize to actin structures involved in migration such as lamellipodia, and in extracellular matrix degradation and cell invasion like invadosomes (*14, 16*).

Paralogous genes, that reflect the high degree of redundancy in the human genome, are ancestrally duplicated genes that frequently retain, at least partially, overlapping functions. Because of the resulting buffering effect, paralogous genes are less likely to be essential for cell growth than non-paralogous ("singleton") genes. Nevertheless, two paralogs may exhibit synthetic lethality where removal of both is deleterious but individual loss is tolerated. Since cancer cell genomes commonly harbor deletions and inactivating mutations in paralogs, synthetic lethality-type interactions between paralogs provide new possibilities for anti-cancer therapy (*17–19*). In this study, we focus on the interplay between p190A and p190B in endometrial cancer and to investigate these paralogs as potential targets. Specifically, we address the p190 paralog dependency and interaction in endometrial cancer.

## RESULTS

### Expression of both *ARHGAP35* and *ARHGAP5* genes is altered in endometrial carcinomas

The TCGA PanCancer Atlas regroups 529 samples of Uterine Corpus Endometrial Carcinomas (UCEC), representing all three tumor types: endometrioid, serous and mixed serous/endometrioid carcinomas **(Figure 1A)**. Analysis of these data using the open-source software CBioPortal for Cancer Genomics (http://www.cbioportal.org/) revealed that somatic mutations, either missense or truncating mutations, are observed at a frequency of about 21% and 10% for respectively *ARHGAP35* and *ARHGAP5* genes in endometrial carcinomas **(Figure 1B)**. Mutations, that include missense, nonsense and in-frame deletions, were identified all along the coding sequence, without hot spots **(Figure 1C)**. We further analyzed the co-occurrence of *ARHGAP35* or *ARHGAP5* mutations with mutations in the twelve most commonly mutated genes in UCEC. We confirmed the co-occurrence of *ARHGAP35* and *POLE* mutations as previously reported (*8*). Interestingly, we observed a strong co-occurrence of mutations in both p190 paralogs in the TCGA cohort **(Figure 1D)**, leading consequently to a strong co-occurrence of *ARHGAP5* and *POLE* mutations. In terms of potential prognostic value, we observed that tumors bearing co-alteration of *ARHGAP35* and *ARHGAP5* genes have a better prognosis than those with only one altered gene or no alteration at all **(Figure 1E)**.

**Figure 1.**
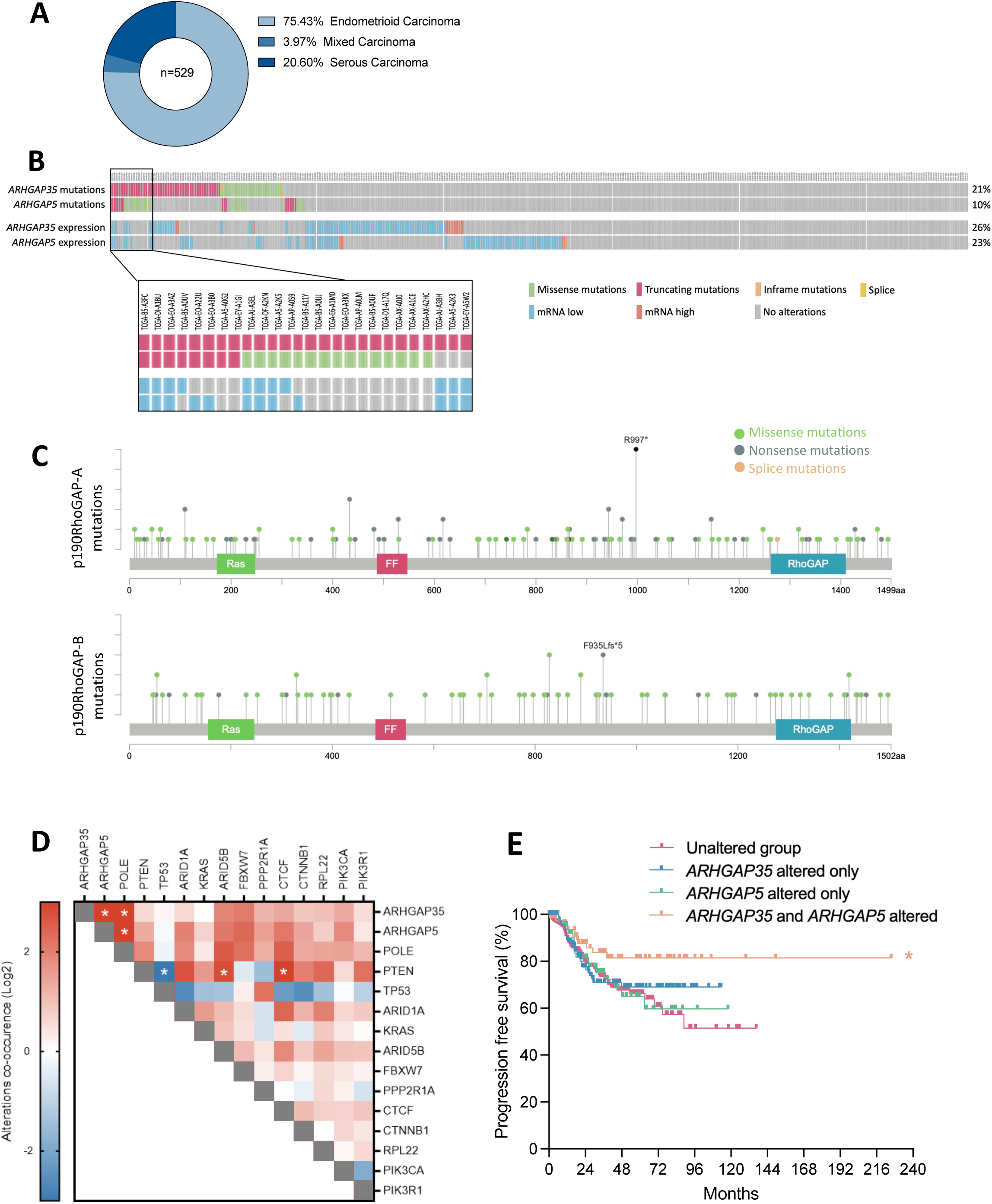
Alterations in *ARHGAP35* and *ARHGAP5* genes in Uterine Corpus Endometrial Carcinoma (TCGA, Pan Cancer Atlas) (A) Distribution of the UCEC tumor subtypes from **TCGA,** cohort (n=529). (B) Representation of *ARHGAP35* and *ARHGAP5* alterations (ie. mutations and mRNA expression) across 529 UCEC cases from TCGA. (C) Lollipop plots showing the repartition of somatic mutations identified in UCEC on p190A (up) and p190B (down) proteins. (D) Co-occurrence of *ARHGAP35* and *ARHGAP5* mutations with 13 most commonly mutated genes in UCEC. Log2 Odds Ratio quantifies how strongly the presence or absence of alterations in A are associated with the presence or absence of alterations in B in the selected samples. OR = (Neither * Both) / (A Not B * B Not A). *log2ratio>3. (E) Progression free survival curve of UCEC cases altered for *ARHGAP35*, *ARHGAP5* or both compared to unaltered group. *p<0.05. All data were extracted from the open-source software cBioPortal for Cancer Genomics (http://www.cbioportal.org/).

The impact of *ARHGAP35* mutations was recently explored for p190A GAP activity, demonstrating either loss-of-functions, gain-of-functions or no functional impact (*9*). Thus, at this stage, it is difficult to anticipate the outcome for cancer cells. Nevertheless, regardless of mutations, we found that mRNA expression of both genes was altered in a large number of endometrial tumors. Indeed, in comparison to normal samples, *ARHGAP35* and *ARHGAP5* mRNA expressions were significantly downregulated in endometrial carcinomas of the TCGA cohort **(Figure 2A)**. Interestingly, we further observed a positive correlation between both paralog expressions **(Figure 2B)**.

**Figure 2.**
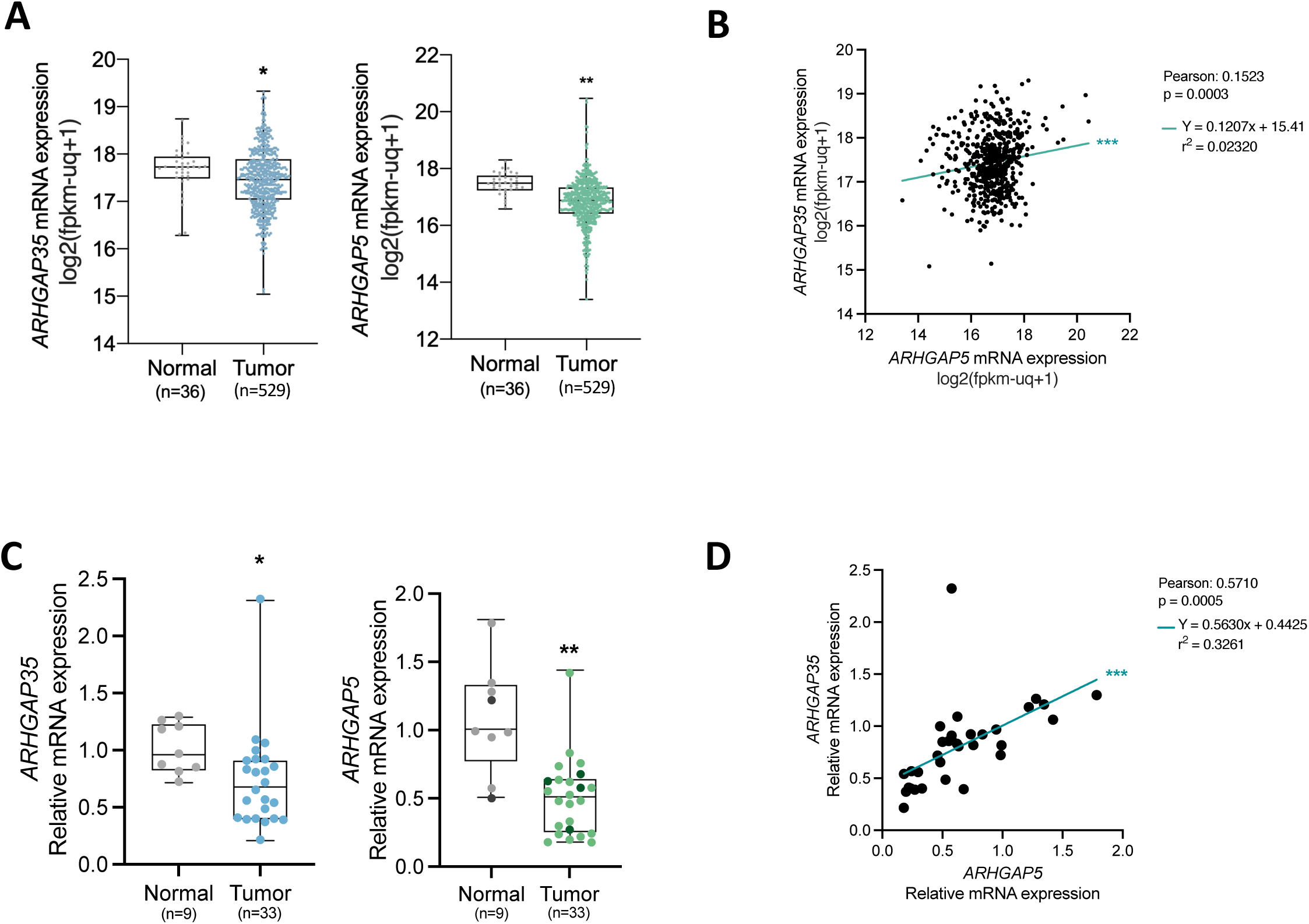
Correlation of mRNA level between *ARHGAP35* and *ARHGAP5* in endometrial tumors. (A) mRNA expression of *ARHGAP35* (left) and A*RHGAP5* (right) between endometrial tumors and healthy tissues in the TCGA cohort. (B) Correlation between *ARHGAP35* and *ARHGAP5* mRNA expression from the TCGA cohort. (C) mRNA expression of *ARHGAP35* (left) and *ARHGAP5* (right) quantified by qPCR in endometrial patient samples. (D) Correlation between *ARHGAP35* and *ARHGAP5* mRNA expression, shown in C. (C, D) Represented values are Fold Change relative to mean of normal tissues. A, C were analyzed by t tests, *p<0.05, **p<0.01, ***p<0.001.

We further explored their expression in a cohort of 33 samples (24 from tumoral regions and 9 from non-tumoral regions) from 16 patients (described in **Table S1**), where we quantified both gene expressions by qRT-PCR on mRNA extracted from frozen samples. Our data confirmed data from TCGA, demonstrating that *ARHGAP35* and *ARHGAP5* gene expressions were significantly downregulated in tumor samples when compared to normal uterine tissues **(Figure 2C)**. Again, data analysis shows a striking positive correlation between *ARHGAP35* and *ARHGAP5* mRNA expressions **(Figure 2D)**.

Finally, we explored the expression of *ARHGAP35* and *ARHGAP5* in endometrial cancer cell lines **(Figure 3)**. HeLa cells and hepatocellular carcinoma Huh7 cells, in which the *ARHGAP5* gene was found amplified (*20*), were used as control. We found that both paralogs were co-expressed in all tested endometrial cell lines at the mRNA (**Figures 3A-B**) and protein (**Figures 3C-D**) levels. We noticed that mRNA and protein levels were not strictly correlated, suggesting post-transcriptional and/or post-translational regulations. Moreover, expression of p190A and p190B were independent on the mutational status of the encoding gene (**Table S2**). Indeed, based on data extracted from the Cancer Cell Line Encyclopedia (CCLE) database (Broad Institute), HEC-1-A and HEC-1-B cell lines are bearing mutations on *ARHGAP35* gene, whereas AN3CA, Ishikawa, KLE and RL95-2 cells lines are mutated on *ARHGAP5* gene.

**Figure 3.**
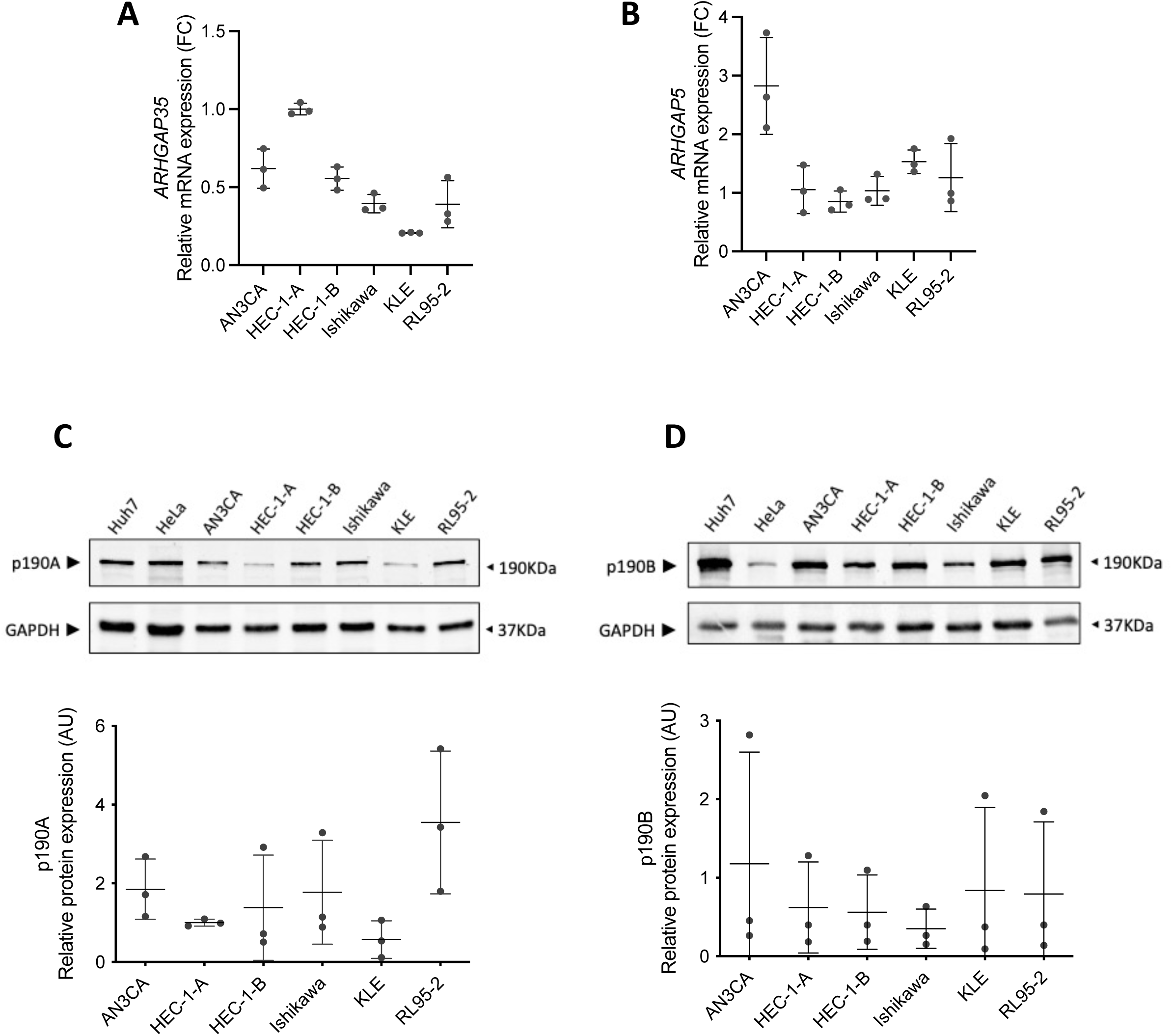
Expression of *ARHGAP35*/p190A and *ARHGAP5*/p190B in endometrial cancer cell lines. (A, B) mRNA expression of *ARHGAP35* (A) and *ARHGAP5* (B) in 6 endometrial cell lines. Fold Change values were calculated after HEC-1-A ΔΔCt mean. mRNA expression is presented relative to expression in HEC1A cells. (C-D) Western blots showing p190A and p190B protein expression in 6 endometrial cell lines. Hepatocellular carcinoma Huh7 cells and HeLa cells were used as control cell lines. Note that as expected p190B is amplified in Huh7 cells. Quantification of p190A (C) and p190B (D) protein expression is presented relative to HEC-1-A cells. AU: arbitrary unit.

### p190A and p190B protein alterations impact similar pathways in endometrial cancer cells

In order to gain further insight into p190A and p190B involvement in endometrial cancer, we generated *ARHGAP35* or *ARHGAP5* knockout (KO) cells and explored their phenotype and behavior. To generate p190A-KO and p190B-KO cell lines, we used CRISPR/Cas9 method with specific sgRNAs in HEC-1-A endometrial cells. We chose to transfect cells with ribonucleoprotein complexes containing sgRNA and Cas9 protein, to allow only a short-term expression of the nuclease in the cells. As shown in Supplementary Figure 1, p190A or p190B protein expression was completely silenced in several HEC-1-A cell clones. Three clones of each were chosen for their biallelic sequence alteration (**Figure S1**). Two KO clones of each paralog, namely KO A-1 and KO A-2 (corresponding respectively to clones 9 and 11) for p190A-KO and KO B-1 and KO B-2 (corresponding respectively to clones 7 and 9) for p190B-KO, were further used for phenotyping (**Figure 4**). Interestingly, we found that removal of one paralog did not alter the expression of the other one **(Figure 4A)**. As p190 proteins are modulators of Rho family members, we first analyzed the impact of each KO on actin cytoskeleton organization. WT and p190-KO HEC-1-A cells were stained with fluorescent phalloidin and analyzed by confocal microscopy using the same laser intensities. Consistent with a loss of Rho GAP activity, we found a striking and significant increase in F-actin staining in both p190A- and p190B-KO cells, when compared to WT cells **(Figure 4B-C)**. A closer analysis of the F-actin organization revealed an unusual remodeling into geodesic F-actin structures in both p190A- and p190B-KO cells **(Figure 4D)**. These structures that show a regular polygonal network of actin fibers, were reminiscent of the, so-called, cross-linked actin networks (CLANs), mainly described in human trabecular meshwork (HTM) cells (*21, 22*). Respectively, 26% and 21% of p190A- and p190B-KO cells displayed CLANs compared to 6% for parental cells **(Figure 4E)**. We found that alpha-actinin co-localized with each F-actin dot within the networks in p190-KO HEC-1-A cells, as described in HTM cells (*22*) **(Figure S2)**, confirming that those p190-KO-induced structures are related to CLANs. To analyze whether CLANs were dependent on RhoA/ROCK pathway, which is expected to be over-activated upon p190A or p190B silencing, we treated p190-KO clones with the Y-27632 inhibitor, targeting Rho-kinase ROCK **(Figure S3)**. We observed a significant decrease in the number of CLANs upon ROCK inhibition, demonstrating that p190-KO-induced CLANS were generated through an increase of RhoA/ROCK activity in HEC-1-A cells.

**Figure 4.**
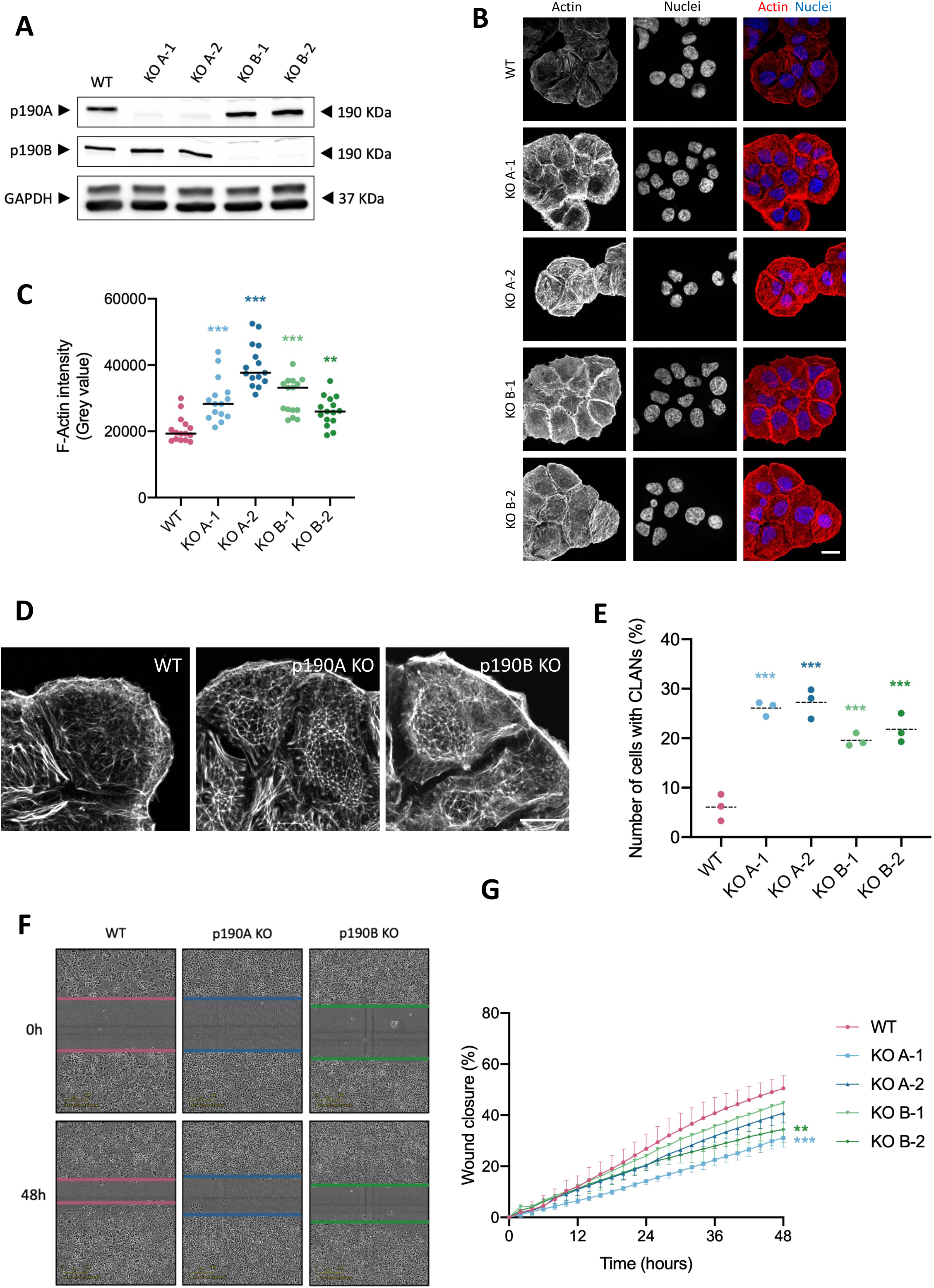
Phenotypical and functional impact of p190A and p190B Knockout in HEC-1-A cells. (A) Western Blots show the absence of expression of p190A and p190B in HEC-1-A KO cells compared to WT cells. KO A-1 and KO A-2 are 2 independent KO clones for p190A; KO B-1 and KO B-2 are 2 independent KO clones for p190B. GAPDH was used as loading control. (B) HEC-1-A WT and KO cells were labelled with phalloidin to stain F-actin (red) and DAPI to stain nuclei (blue). KO and WT cells were imaged with the same exposure time. Scale bar: 10 μm. (C) Quantification of F-Actin intensity in phalloidin-labelled WT and KO HEC-1-A cells. Mean fluorescence intensity was measured across the cells, in triplicate, with n=5 cells each time representing 15 measured points on the figure. The black line represents mean. (D) WT and KO cells are stained with phalloidin showing the organization of the actin cytoskeleton. CLANs structures were observed in p190-KO cells. Representing images of p190A and p190B KO cells are respectively KO A-1 and KO B-1 cells. Scale bars: 10 μm. (E) Quantification of the number of cells containing CLANs. Quantification was made in triplicate with n>100 cells for each condition; mean is represented by the dashed line. (F) Representative images of Wound healing assay in WT or p190-KO HEC-1-A cells at t=0h and t=48h. Images of p190A and p190B KO cells are respectively KO A-1 and KO B-1 cells. (G) Cell migration represented by wound closure across time. (C, E, G) Results were analyzed by 2-way ANOVA, **p<0.01, ***p<0.001 when compared to WT condition.

With regard to this atypical reorganization of the actin cytoskeleton, which suggests a potential impact on cell motility, we further analyzed KO cell migration using wound-healing assay **(Figure 4F-G)**. We found that migration is slightly reduced for p190A-KO cells, and significantly decreased for p190B-KO cells when compared to WT HEC-1-A cells, demonstrating that both p190 paralog expressions are required for proper endometrial tumor cell migration.

Altogether, this analysis, done in parallel for p190A- and p190B-KO clones demonstrated that both paralogs are involved at the same extend in actin remodeling, RhoA/ROCK pathway activation and cell migration in HEC-1-A cells. Moreover, these data suggest that, in endometrial cells, loss of p190A is not compensated by p190B and vice-versa.

### Targeting both p190A and p190B reveals vulnerabilities in endometrial cancer cells

In order to better describe and compare alterations upon p190A or p190B loss, we performed a global proteomic analysis of all 6 clones generated by CRISPR (p190A-KO A-1, A-2 and A-3; p190B-KO B-1, B-2 and B-3) using mass spectrometry. A total of 4159 and proteins were commonly identified in p190A- and p190B-KO cells. A total of 536 proteins in p190A-KO HEC-1-A cells and of 279 in p190B-KO cells showed significant altered expression compared with control cells **(Figures 5A and C)**. To investigate the functional association of the differentially expressed proteins, we conducted Ingenuity Pathway Analysis to identify the most significant pathways that were altered in p190A-KO and p190B-KO HEC-1-A cells. Interestingly, even if altered proteins were different, the analysis pointed out similar signaling pathways. Besides the pathways of cellular organization and cellular movement that were related to alteration of the actin cytoskeleton, we found alteration in other pathways such as cell death and survival, cell proliferation and cellular response to therapeutics suggesting that both p190A and p190B may likely interfere with these pathways **(Figures 5B and D)**. Thus, we first focused on the potential role of p190A and p190B in endometrial cell proliferation and cellular response to therapeutics. Cell growth was monitored using the IncuCyte^®^ system by analyzing the occupied area (percentage of confluence) of cells over time. We observed a slight impact of the down-expression of p190A and p190B on HEC-1-A growth rate, with a higher impact upon p190B loss **(Figure 6A)**. We further tested the sensitivity of p190-KO clones towards cisplatin, which remains the most current therapy used for endometrial tumors. Upon treatment with 100 µM of cisplatin, we found that, as WT cells, p190-KO cells are strongly affected in their growth **(Figure S4)**, suggesting that loss of p190A or p190B did not modify HEC-1-A cell response to cisplatin. Thus, we decided next to explore the potential synthetic lethality between the two paralogs. To do so, we used RNA interference to knockdown the second paralog in KO clones, and analyzed HEC-1-A cell growth in these conditions **(Figure 6B-D)**. We showed that removal of p190B in p190A-KO cells led to a strong decrease of cell growth **(Figure 6C)**, whereas removal of p190A in p190B-KO cells led to only a small impact **(Figure 6D)**, reflecting probably a threshold effect of silencing. This experiment of single and double silencing of paralogs was repeated using exclusively RNA interference in HEC-1-A and RL95-2 cells **(Figure 6E)**. We found that removal of both paralogs was deleterious in HEC-1-A and RL95-2 cells whereas individual loss was tolerated in RL95-2 cells. To define more precisely the synthetic lethality interactions between these two paralogs, we assessed cell viability in WT versus p190A and/or p190B KD cells. To do so, we silenced one or both paralogs by using RNA interference in HEC-1-A cells. As shown on **Figure 6F**, loss of only one form impaired cell viability but loss of the two paralogs leads to a dramatic decrease in cell viability, which confirms the cell death and survival pathway identified above. Altogether, these data demonstrate that p190A and p190B proteins harbor synthetic lethality-type interactions that may represent a potential target for therapy against endometrial cancer.

**Figure 5.**
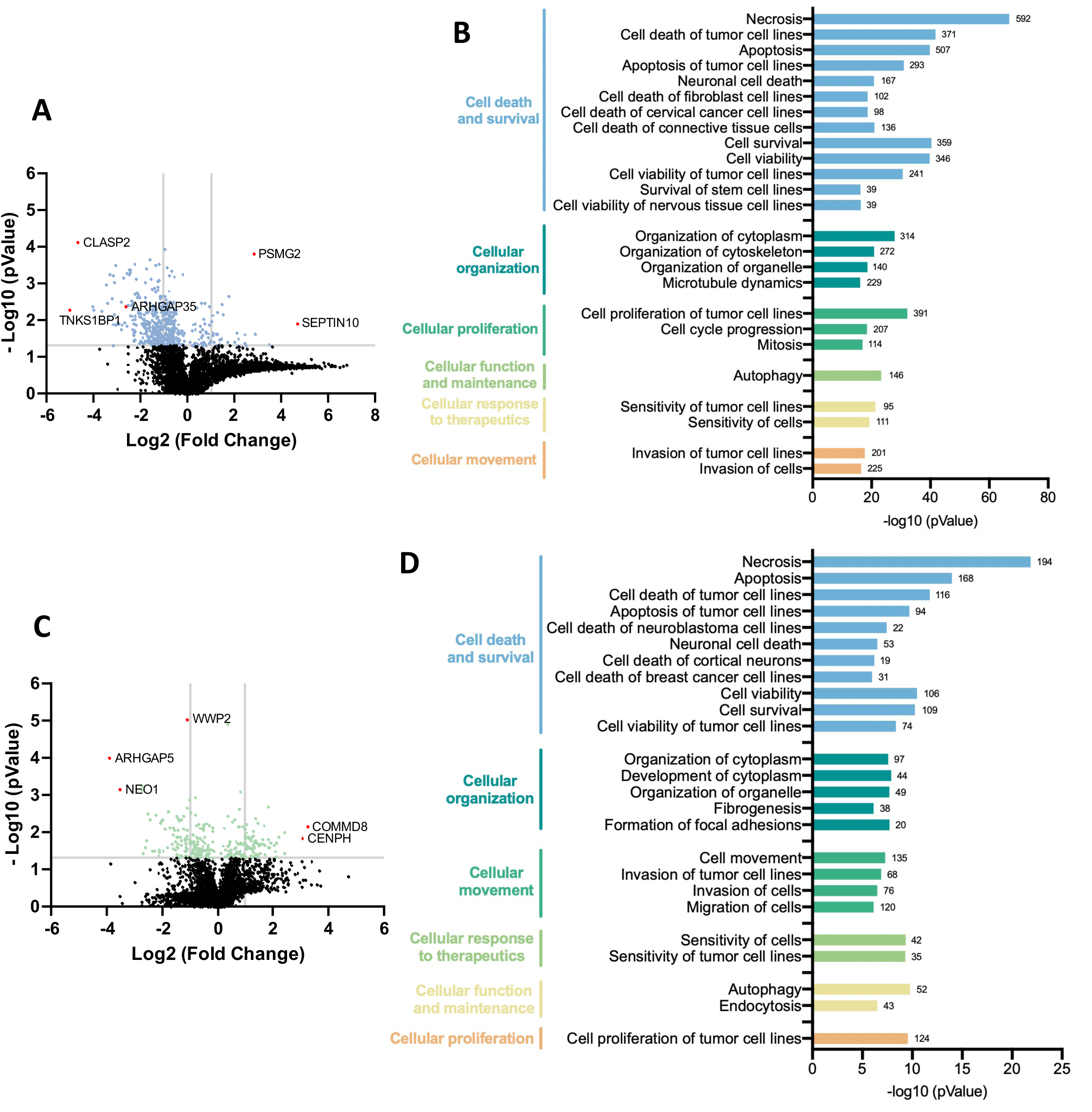
Global proteomic analysis of p190A and p190B KO cells compared with WT HEC-1-A. Three clones of each KO (KO A-1, A-2 and A-3 for p190A; KO B-1, B-2 and B-3 for p190B) were analysed and combined. (A,C) Volcano Plot highlighting the top up- and downregulated proteins in p190A KO (A) and in p190B KO (C) cells compared to WT HEC-1-A. Fold changes were calculated for each protein after the WT value. Most deregulated proteins are indicated (red dots). (B, D) IPA enrichment analysis of the top deregulated cellular functions in p190A KO (B) and in p190B KO (D) HEC-1-A cells. The number of deregulated proteins in each category is indicated at the end of bars.

**Figure 6.**
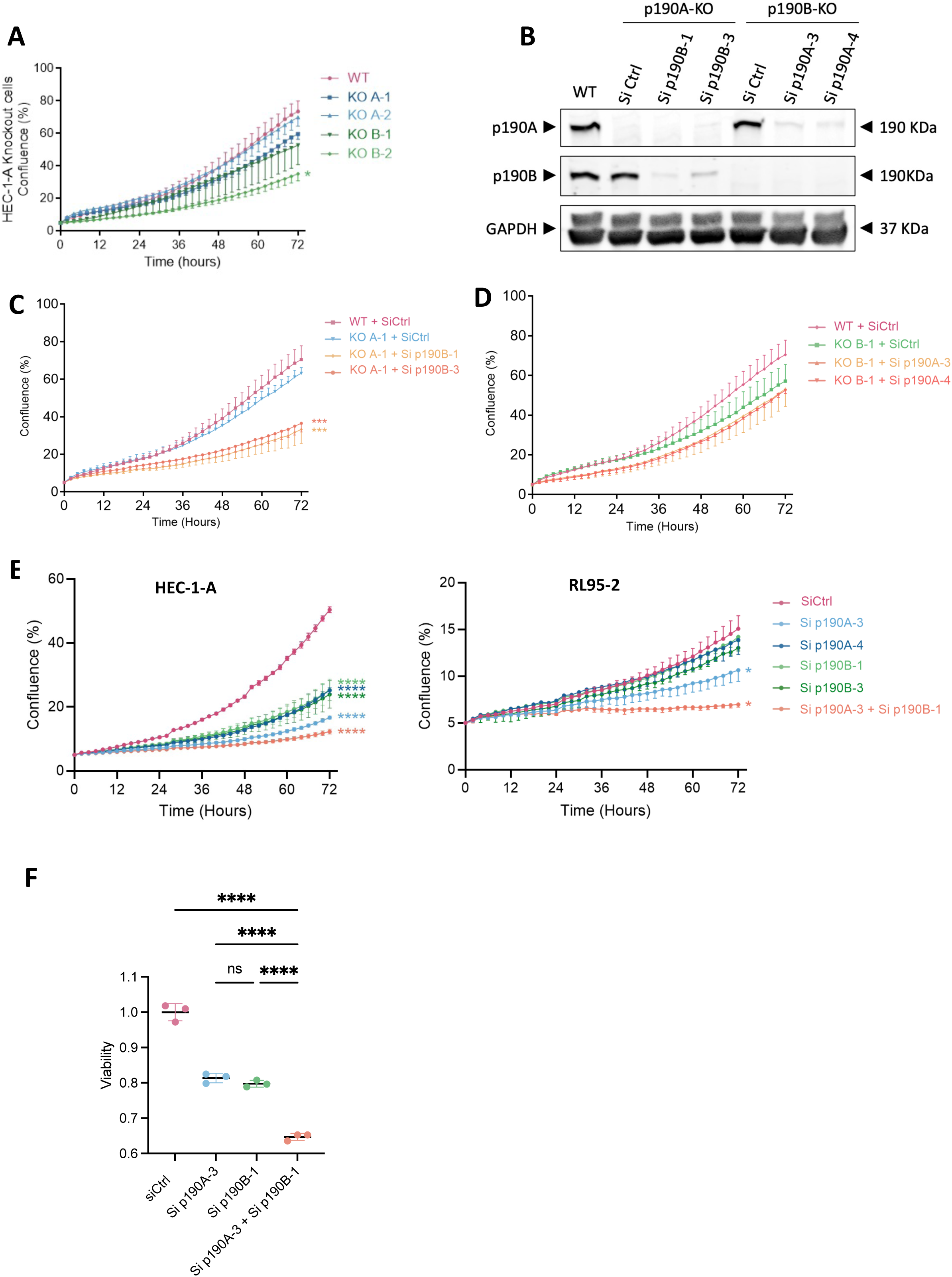
p190RhoGAPs expression inhibition impacts cellular proliferation and cell viability of endometrial cell lines. (A) Proliferation of WT vs p190A- and p190B-KO HEC-1-A cells (B) Western Blots showing the extinction of p190A and p190B expression using siRNA in KO cells leading to a double KD/KO of both paralogs. Sip190B-1 and sip190B-3 are two independent siRNAs targeting p190B expression, same for p190A with sip190A-3 and sip190A-4. (C, D) Proliferation of KD/KO cells analyzed in p190B siRNA transfected HEC-1-A p190A-KO cells (C) and in p190A siRNA transfected HEC-1-A p190B-KO cells (D). (E) Proliferation of HEC-1-A (left-hand graph), and RL95-2 (right-hand graph) endometrial cell lines upon expression inhibition of p190A and/or p190B using siRNAs. (F) Cell viability was assessed in double siRNA transfected HEC-1-A cells. (A-F) Figures were analyzed by 2-way ANOVA, *p<0.05, ***p<0.001, ****p<0.0001 when compared to WT conditions.

## DISCUSSION

*ARHGAP3*5/p190A and A*RHGAP5*/p190B are paralogs ubiquitously expressed in tissues. Herein we addressed their expression and function in endometrial cancer, in which *ARHGAP35* was reported to be frequently mutated and described as a cancer gene (*5, 6*). In this study, we revealed that *ARHGAP5* gene is also mutated in endometrial tumors, even if at a lower frequency than *ARHGAP35* gene. Interestingly, a co-occurrence of *ARHGAP35* and *ARHGAP5* mutations occurs in endometrial cancer, reflecting paralog’s interplay and suggesting a common mechanism responsible for mutation appearance. We also found a co-occurrence of *ARHGAP35* or *ARHGAP5* and *POLE* mutations in endometrial cancer. *POLE* gene encodes the catalytic subunit of the DNA polymerase Epsilon, and *POLE* mutations were associated with the disruption of the exonuclease activity of the polymerase required for its proofreading function. Whether *ARHGAP35* and *ARHGAP5* mutations are a consequence of this POLE dysfunction remains to be explored. However, some insight comes from normal endometrial tissues. Indeed, endometrium was described as one of the tissues that surprisingly displays frequent mutations in cancer genes (*23, 24*). *ARHGAP35* is the most frequently mutated gene in normal uterine endometrial epithelium (fraction of mutated samples of 0.34), even more frequently than in the corresponding endometrial cancer (*24*). On the other hand, *POLE* mutations remain at a very low frequency in normal endometrium, suggesting that emergence of *ARHGAP35* mutations is not due to DNA Polymerase Epsilon dysfunction. On the contrary, *POLE* mutations shown to increase genomic instability may lead to increased formation of neoantigens that may in turn favor an early immune elimination of the transformed cells. This hypothesis, which needs further exploration, would give a potential explanation to the diminution of frequency of *ARHGAP35* mutations in endometrial cancer compared to normal endometrium.

Regardless of mutations, we further demonstrated that *ARHGAP35* and *ARHGAP5* expressions are significantly downregulated in tumors when compared to non-tumoral tissues. Moreover, we described a positive correlation of expression between *ARHGAP35* and *ARHGAP5* genes, both in the TCGA cohort and in our own cohort. These results suggest that both paralogs may share common regulatory mechanisms, as already described for close paralogs (*25*). Thus, *ARHGAP35* and *ARHGAP5* may share common transcription factor binding sites in their promoters or enhancer elements active in endometrial cancers. However, it remains to be explored if this correlation in gene expression is specific of endometrial cancers. Even though *ARHGAP35* and *ARHGAP5* genes are not located on the same chromosome, evolutionary constraints in functional genome organization such as three-dimensional chromatin architecture may be responsible for such concerted expression (*26*).

In order to mimic the low levels of proteins encoded by *ARHGAP35* and *ARHGAP5* genes in endometrial cancers, we generated and characterized KO cells. We found that KO of p190A or p190B led to an important increase of actin filaments in cells. This phenotype is consistent with the fact that p190 proteins are negative regulators of Rho GTPases. Loss of these negative regulators favors G- to F-actin polymerization. In various cells such as fibroblasts, p190A or p190B was associated to the regulation of actin stress fibers formation through the control of RhoA activity (*27–29*). In contrast, we found that p190A- or p190B-KO cells displayed an atypical remodeling of the actin cytoskeleton, showing geodesic F-actin structures reminiscent of CLANs, observed mainly in HTM cells. HTM cells are endothelial-like cells in a sponge-like connective tissue located near the front of the eye and are responsible for regulating eye pressure. CLANs are thought to play a role in resistance of cells to increased intraocular pressure that is associated to glaucoma, by creating stiffer cells and tissue (*21*), and they were also associated to the pathology inducing trabecular meshwork dysfunction (*30*). The presence of CLANs in endometrial cancer cells upon p190A- or p190B-KO suggest that those cells might adapt differently to their environment, and increase endometrium stiffness. We further demonstrated that these actin networks are highly dependent on the RhoA/ROCK pathway and deleterious for cell migration. Whether CLANs are involved in the development of endometrial pathologies such as cancer is of interest and remains to be explored.

Global analysis of altered proteins upon p190A- or p190B-KO in HEC-1-A cells revealed that both proteins have overlapping functions. Indeed, similar cellular functions were altered upon silencing of each paralog, opening the path to create vulnerabilities in endometrial cancer cells. We thus explored their redundancy into cell growth assays and into cell viability. We found that removal of one paralog tends to decrease endometrial cell growth as well as cell viability, and that removal of both has a stronger impact. Until a certain threshold, reduced levels of p190A or/and p190B may remain tolerated in cancer cells due to a buffering effect between both proteins. However, below this threshold that we mimic using co-knockdown of p190A and p190B, paralogs became essential and their loss is deleterious. Thus, these finding demonstrated the tight interplay between p190A and p190B proteins and open a new avenue to design novel strategy to target endometrial cancer cells.

## EXPERIMENTAL PROCEDURES

### Cell lines

HEC-1-A cell line was cultured in McCoy’s 5A medium. HEC-1-B, AN3CA, Ishikawa cell lines were cultured in MEM complete medium. KLE and RL95-2 cell lines were cultured in DMEM:F-12 Medium. All media were supplemented with 10% FCS (Eurobio). Cells were maintained at 37°C in a 5% CO_2_ humidified atmosphere. HEC-1-A, HEC-1-B, KLE and RL95-2 cell lines were purchased from American Type Culture Collection (ATCC). AN3CA and Ishikawa cell lines were purchased from Cell Lines Services and Sigma Aldrich, respectively. Hepatocellular carcinoma Huh7 cells, in which the *ARHGAP5* gene was found amplified (*20*), and HeLa cells were cultured as described previously (*13*) (*31*). Cell lines were confirmed for the absence of mycoplasma by PCR.

### RT-qPCR

Frozen fractions of endometrial samples were crushed manually using a liquid nitrogen-cold mortar. mRNAs from both endometrial tissues and cultured cells were extracted using the Nucleospin RNA^®^ kit from Macherey Nagel according to the manufacturer’s instructions. cDNA was synthetized from 1 µg of total RNA with Maxima Reverse Transcriptase (Fermentas). 30 ng of cDNA were then subjected to PCR amplification on an RT-qPCR system using the CFX96 Real Time PCR detection system (Biorad). The SYBR^®^ Green SuperMix for iQTM (Quanta Biosciences, Inc.) was used with the following PCR amplification cycles: initial denaturation (95 °C for 10 min), followed by 40 cycles of denaturation (95 °C for 15 s) and extension (60 °C for 1 min). Gene expression results were first normalized to an internal control with 18 S ribosomal RNA. Relative expression levels were calculated using the comparative (2-ΔΔCT) method.

### Antibodies and reagents

Mouse monoclonal anti-p190A (clone 30; ref#610149) and anti-p190B (clone 54; ref#611612) antibodies were purchased from DB Transduction Laboratories. Mouse monoclonal anti-p190A (clone D2D6; ref#R3150) and anti-ɑ-actinin (clone EA-53; ref#A7811) antibodies were purchased from Sigma Aldrich. Polyclonal anti-p190B antibody (ref#2562S) was purchased from Cell Signaling Technology. Rabbit (FL-335; ref#sc-25778) and mouse (D-6; ref#sc-166545) anti-GAPDH antibodies were obtained from Santa Cruz Biotechnology. Anti-mouse IRDYE680 and anti-rabbit IRDYE800 secondary antibodies were obtained from Eurobio Scientific. FluoProbes 488-phalloidin, 488-labeled secondary antibodies and 547-labeled secondary antibodies were purchased from Interchim. Y-27632 (Sigma Aldrich) was used at 10 μM for 24 hours. Cisplatin (MedChemTronica) was used at 100 µM for 72 hours.

### Transfections

SiRNAs were transfected using Lipofectamine RNAiMAX (ThermoFisher Scientific) according to the manufacturer’s protocol. SiRNAs targeting p190A were 5’-GCAGAAGUUGCAUGCCCUUAA-3’ and 5’- GAACAGCGAUUUAAAGCAUUU-3’ for respectively sip190A-3 and sip190A-4. SiRNAs targeting p190B were 5’-AACGTGCAGCTGCATCTAAAT-3’ and 5’-AATGAGAAGCATATCTGGTTA-3’ for respectively sip190B-1 and sip190B-3. The control siRNA corresponds to All Stars Negative control from Qiagen.

### Western blot analysis

Cells were scraped off on ice and homogenized in Ripa lysis buffer (50 mM Tris HCl, pH 7.5, 150 mM NaCl, 0.5% sodium deoxycholate, 5mM EDTA, 0.1% SDS) with protease and phosphatase inhibitors. Cell lysates were cleared of cellular debris and nuclei by a 10,000 g centrifugation step for 15 min. Lysates were denatured with Laemmli loading buffer containing 2.5% 2-β-mercaptoethanol, analyzed by SDS-PAGE, and blotted onto nitrocellulose membranes. Blots were incubated overnight at 4°C with primary antibodies, and then incubated with infrared fluorescent dye-conjugated secondary antibodies (Eurobio). Signals were visualized with the Chemidoc infrared imaging system (Bio-Rad) and analyzed using ImageJ image analysis software.

### Generation of CRISPR/Cas9-mediated KO cell lines

The Clustered Regularly Interspaced Short Palindromic Repeats (CRISPR) system was used as a genome editing method to delete *ARHGAP35* or *ARHGAP5* gene in HEC-1-A cells. For each gene, two gRNAs were designed with CRISPOR algorithm (https://crispor.gi.ucs.edu (*32*)): *ARHGAP35*-gRNA#1 (5’- CAGCACTCGGGCGCACGAAG-3’), *ARHGAP35*-gRNA#2 (5’-TGCAGGGCCGTGCTTCGATG-3’), *ARHGAP5*- gRNA#1 (5’-CTATACTGATGGTATAGGAT-3’), *ARHGAP5*-gRNA#2 (5’-TCAGTATAGTTGGACTCTCT-3’). Alt-R^®^-crRNA corresponding to the target sequences were purchased from IDT (Coralville, IA, USA). Alt-R^®^ S.p-Cas9HIFI (4.5 pmol corresponding to 750 ng, IDT) was mixed with a 1.2 excess molar ratio of two-part gRNA (Alt-R^®^-crRNA +Alt-R^®^-tracrRNA) reconstituted following the supplier’s recommendations. Ribonucleoprotein complexes were transfected in 40,000 HEC-1-A cells using Lipofectamine CRISPRMAX reagent (Thermo Fischer Scientific). At 2-3 days post transfection, a sample of cells was lysed and used as PCR template using Phire Tissue Direct PCR Master Mix (Thermo Fisher Scientific). PCR amplification with subsequent Sanger sequencing (Genewiz) of the targeted loci was performed according to the supplier’s recommendations with primers 5’-GCGATGAGAGGTGGTGAACA-3’, 5’- CAGCTGATGCAAGCTTGGTC-3’ and 5’-GCTTTCCGTCTGGCATTTGT-3’ for *ARHGAP35* gene and primers 5’-TATGGTGCTTTGTTGTAAACATCT-3’ and 5’- TCCTCCAAAGTCAATGGTGCT-3’ for *ARHGAP5* gene. Sanger chromatograms analysis using the ICE (https://ice.synthego.com (*33*)) and Tide (https://tide.nki.nl (*34*)) algorithms has confirmed a better modification efficiency with gRNA#1 for both genes. gRNA#1 transfected cells were further seeded in a 96-well plate by limited dilution to isolate monoclonal cell lines which were then genotyped by PCR and Sanger sequencing to identify those carrying mutations that create a frameshift leading to a gene Knock-Out. KO clonal cell lines were further validated by Western-blot. HEC-1-A KO cell cultures were maintained and handled under the same conditions reported for the WT.

### Immunofluorescence and confocal imaging

Cells were seeded on glass coverslips coated with fibronectin (Thermo Fisher Scientific), for 48 hours. They were fixed with 4% PFA for 10 min at RT and permeabilized with 0.2% Triton X-100 for 5 min before incubation with various antibodies, as previously described (*35*). After staining, coverslips were mounted on slides with Fluoromount G mounting medium. Cells were imaged using an SP5 confocal microscope (Leica Biosystems) using a 63×/NA 1.4 Plan Neofluor objective lens and the LAS-AF-Lite 2.4.1 acquisition software (Leica Biosystems). To prevent contamination among fluorochromes, each channel was imaged sequentially using the multitrack recording module before merging. Images were processed using LAS-AF-Lite 2.4.1 (Leica Biosystems) or ImageJ software (National Institutes of Health). Phalloidin staining intensity was analyzed using ImageJ software, by measuring grey values across cells using images captured with the same laser intensities.

### IncuCyte^®^ assays

Proliferation assays: Cells were seeded in 96-well plates (3.10^3^ cells per well) and then monitored by the IncuCyte^®^ Zoom videomicroscope (Essen Bioscience). Cell confluence and quantification was measured using the IncuCyte^®^ imaging system. For Cisplatin experiments, Cisplatin was added at 100 µM. Migration assays: Cells were seeded in appropriate 96-well plates (5.10^4^ cells per well) (Essen Bioscience). After seeding, scratch was made using the Wound Maker (Essen Bioscience), and wound healing was monitored by the IncuCyte^®^ Zoom videomicroscope. Quantification was measured using the IncuCyte^®^ imaging system.

### Mass spectrometry-based label-free quantitative proteomics

Cell lysis was performed in RIPA Buffer. 10 µg of proteins were loaded on a 10% acrylamide SDS-PAGE gel and proteins were visualized by Colloidal Blue staining. Sample preparation, protein digestion by trypsin and LC-MS parameters used for nanoLC-MS/MS analysis nanospray Orbitrap Fusion™ Lumos™ Tribrid™ Mass Spectrometer (Thermo Fisher Scientific, California, USA) were previously described (*36*).

### Mass spectrometry data analysis

Protein identification and label-free quantification (LFQ) were done in Proteome Discoverer 3.1. The CHIMERYS node using the prediction model inferys_3.0.0 fragmentation was used to identify proteins in batch mode by searching against the UniProt *Homo sapiens* database (82233 entries, released June 2024). Two missed enzyme cleavages were allowed for trypsin. Peptide lengths of 7–30 amino acids, a maximum of 3 modifications, charges of 2–4, and 0.6 ppm for fragment mass tolerance were set. Oxidation (M) and carbamidomethyl (C) were respectively searched as dynamic and static modifications by the CHIMERYS software. Only “high confidence” peptides were retained corresponding to a 1% false discovery rate at the peptide level. Minora feature detector node (LFQ) was used along with the feature mapper and precursor ions quantifier. The normalization parameters were selected as follows: (1) Unique peptides, (2) Precursor abundance based on intensity, (3) Normalization mode: total peptide amount, (4) Protein abundance calculation: summed abundances, (5) Protein ratio calculation: pairwise ratio based and (6) Missing values were replaced with random values sampled from the lower 5% of detected values. Quantitative data were considered for master proteins, quantified by a minimum of 2 unique peptides, a fold changes above 2 and a statistical *p-value* adjusted using *Benjamini-Hochberg* correction for the FDR lower than 0.05. The mass spectrometry proteomics data have been deposited to the ProteomeXchange Consortium via the PRIDE (*37*) partner repository with the dataset identifier PXD067294.

### Cell viability assay

Cells pre-treated for 96 hours with siRNAs were seeded in opaque-walled 96-well plates (10000 cells per well) and incubated 24 h at 37 °C. Cells were then lysed and cell viability was assayed using CellTiter-Glo^®^ Luminescent Cell Viability Assay (Promega).

### Statistical analysis

Statistical analysis was performed with Prism software (GraphPad Software) and data are presented as mean ± SEM of at least three independent experiments. Comparisons between two groups were analyzed by t test, while comparisons between several groups were analyzed by 2-way Significance was accepted for values where *, P < 0.05; **, P < 0.01; ***, P < 0.001; and ****, P < 0.0001.

## Supporting information

This article contains supporting information.

## Acknowledgments

We are grateful to Sylvaine Di Tommaso (OncoProt platform, TBMCore, Univ. Bordeaux, CNRS UAR 3427, Inserm US05) for her help in the proteomic approach. Microscopy was done in the Bordeaux Imaging Center, a service unit of the CNRS-INSERM and Bordeaux University, member of the national infrastructure France Bio Imaging. qRT-PCR experiments were performed at the OneCell facility platform (TBMCore, Univ. Bordeaux, CNRS UAR 3427, Inserm US05), CRISPR experiments were done at CRISP’edit facility (TBMCore, Univ. Bordeaux, CNRS UAR 3427, Inserm US05)

## Author contributions

V.M. conceptualization; M.P., C.H., M.CdO, A.A.R., S.C., V.L. and V.M. formal analysis; M. P, C.H., V.N., V.V., A.A.R., V.P-M., J-W. D., B. T. and F.S. methodology; S.C. providing human tumor samples; V.L. and V.M. supervision; V.M. writing original draft; M.P., V.L. and V.M. writing, review and editing; V.M. funding acquisition.

## Data Availability Statement

The data supporting the findings of this study can be found in the article, or available from the corresponding author upon reasonable request.

## Abbreviations

CNV: copy number variation
Cter: Carboxy-terminal
GAP: GTPase-Activating Protein
GBD: GTP-binding domain
GTPase: guanosine triphosphate hydrolase
HTM: Human Trabecular Meshwork
Nter: Amino-terminal
PLS: protrusion localization sequence
Ras: Rat sarcoma
Rho: Ras homolog
WT: wild-type

