## supplementary material for "p190A/*ARHGAP35* and p190B/*ARHGAP5* proteins in endometrial cancer, a novel cancer-relevant paralog interplay"

Supplementary Figure 1 – **Screening of the HEC-1-A clones after applying the CRISPR/Cas9 technique to produce knockout of p190A (A-B) and p190B (C-D).**

(A, C) Western Blots showing the expression or extinction of p190A (A) and p190B (C) proteins in the clones. Protein extract of each clone was analyzed by two independent antibodies against the targeted protein. Three clones of each KO, in which the targeted protein was absent, were chosen for the study. The clones are highlighted in blue for p190A-KO clones and in green for p190B-KO clones. They were renamed KO A-1 (clone#9), KO A-2 (clone #11) and KO A-3 (clone #16) for p190A-KO and KO B-1 (clone#7), KO B-2 (clone #9) and KO B-3 (clone #8) for p190B-KO. (B, D) The predicted sequence of each allele after Sanger chromatogram analysis is shown. The Start codon is highlighted in orange, the altered sequenced is shown in red and the PAM sequence is underlined.

Supplementary Figure 2 – **Alpha-actinin colocalizes with CLANs in HEC-1-A p190 Knockout cells.**

Representative confocal images of HEC-1-A p190A-KO (KO A-1) and HEC-1-A p190B-KO (KO B-1) cells containing CLANs. Phalloidin (red) and α-actinin (green) staining colocalizes in the structures. Nuclei were stained using DAPI (blue). The arrows indicate CLAN nodes. Scale bar: 10 µm.

Supplementary Figure 3 – **CLANs formation is dependent on RhoA/ROCK pathway in p190-KO HEC-1-A cells.** WT, p190A-KO and p190B-KO cells were treated or not with 10 µM of Y-27632 for 24 h. Cells were fixed, stained for F-actin and nuclei using respectively fluorescent phalloidin and DAPI. CLANs were quantified under the microscope. The graph shows the quantification of the presence of CLANs in WT and KO HEC-1-A cells. Results were analyzed by one-way ANOVA, $^{**}p<0.01$, $^{***}p<0.001$.

Supplementary Figure 4 – **p190A/B Knockout did not confer cisplatin resistance to HEC-1-A cells.** Cells were untreated or treated with Cisplatin (100 µM) for 72 hours. Cell proliferation was measured during treatment using the IncuCyte system.

Supplementary Table 1: **Clinical and tumor characteristics of patients used in this study.** Patients are numbered from P1 to P16. Sample information is indicated as "NT(1)/T(3)," where "T" refers to tumoral tissue, "NT" to non-tumoral tissue, and the number in parentheses denotes the number of samples obtained from each patient. AWD, alive with disease; DOD, died of disease; DWD, died with disease; NED, no evidence of disease.

Supplementary Table 2: **Mutational status of endometrial cell lines for *ARHGAP35* and *ARHGAP5* genes.** Data were extracted from the Cancer Cell Line Encyclopedia (CCLE) database (https://sites.broadinstitute.org/ccle/)
