## Supplemental figures for "p190A/*ARHGAP35* and p190B/*ARHGAP5* proteins in endometrial cancer, a novel cancer-relevant paralog interplay"

**A**

#### ① Anti-p190A Ab #1 (Clone 30, BD Transd. Lab)

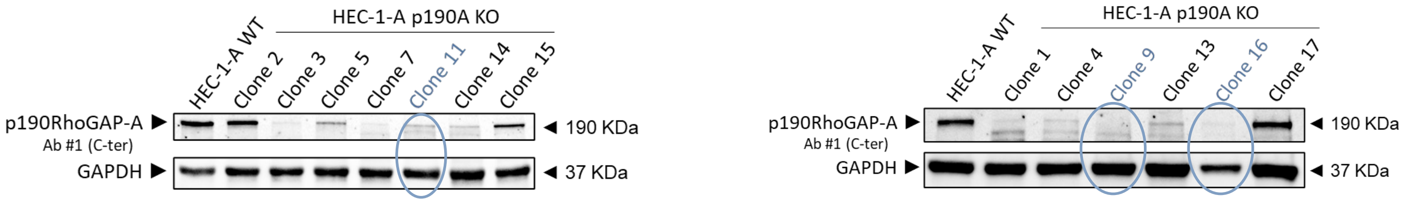


#### ② Anti-p190A Ab #2 (Clone D2D6, Sigma))

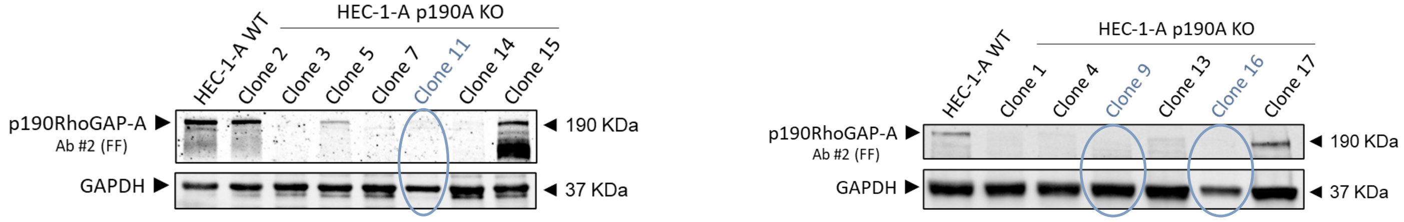


**B**

##### Clone #9 (KO-A#1)

- **Allele n°1: -1 deletion (red)**

--*ATG*ATGATGGCAAGAAAGCAAGATGTCCGAATTCCCACCTACAAC
ATCAGTGTGGTGGGATTATCTGGGACCGAGAAGGAAAAGGGCCAGTGTGGGAT
TGGAAAGTCTTGTTTGTGCAACCGCTT~~**C**~~GTGCGCCCGAGTGCTGACGAGTTTC
ACTTGGACCATACCTCCGTCCTCAGCACCAGTGACTTTGGAGGGCGAGTGGTC
AATAATGACCACTTTCTCTACTGGGGA--

- **Allele n°2: -13 deletion (red)**

--*ATG*ATGATGGCAAGAAAGCAAGATGTCCGAATTCCCACCTACAAC
ATCAGTGTGGTGGGATTATCTGGGACCGAGAAGGAAAAGGGCCAGTGTGGGAT
TGGAAAGTCTTGTTTGTGC~~AACCGCTTCGTGC~~GCCCGAGTGCTGACGAGTTTC
ACTTGGACCATACCTCCGTCCTCAGCACCAGTGACTTTGGAGGGCGAGTGGTC
AATAATGACCACTTTCTCTACTGGGGA–

##### Clone #11 (KO-A#2)

- **Allele n°1: -11 deletion (red)**

--*ATG*ATGATGGCAAGAAAGCAAGATGTCCGAATTCCCACCTACAAC
ATCAGTGTGGTGGGATTATCTGGGACCGAGAAGGAAAAGGGCCAGTGTGGGAT
TGGAAAGTCTTGTTTGTG~~CAACCGCTTCG~~TGCGCCCGAGTGCTGACGAGTTTC
ACTTGGACCATACCTCCGTCCTCAGCACCAGTGACTTTGGAGGGCGAGTGGTC
AATAATGACCACTTTCTCTACTGGGGA--

- **Allele n°2: -13 deletion (red)**

--*ATG*ATGATGGCAAGAAAGCAAGATGTCCGAATTCCCACCTACAAC
ATCAGTGTGGTGGGATTATCTGGGACCGAGAAGGAAAAGGGCCAGTGTGGGAT
TGGAAAGTCTTGTTTGTGCAACCG~~CTTCGTGCGCCCG~~AGTGCTGACGAGTTTC
ACTTGGACCATACCTCCGTCCTCAGCACCAGTGACTTTGGAGGGCGAGTGGTC
AATAATGACCACTTTCTCTACTGGGGA–

##### Clone #16 (KO-A#3)

- **Allele n°1: -13 deletion (red)**

--*ATG*ATGATGGCAAGAAAGCAAGATGTCCGAATTCCCACCTACAAC
ATCAGTGTGGTGGGATTATCTGGGACCGAGAAGGAAAAGGGCCAGTGTGGGAT
TGGAAAGTCTTGTTTGTGCAACCG~~CTTCGTGCGCCCG~~AGTGCTGACGAGTTTC
ACTTGGACCATACCTCCGTCCTCAGCACCAGTGACTTTGGAGGGCGAGTGGTC
AATAATGACCACTTTCTCTACTGGGGA--

- **Allele n°2: -11 deletion (red)**

--*ATG*ATGATGGCAAGAAAGCAAGATGTCCGAATTCCCACCTACAAC
ATCAGTGTGGTGGGATTATCTGGGACCGAGAAGGAAAAGGGCCAGTGTGGGAT
TGGAAAGTCTTGTTTGTG~~CAACCGCTTCG~~TGCGCCCGAGTGCTGACGAGTTTC
ACTTGGACCATACCTCCGTCCTCAGCACCAGTGACTTTGGAGGGCGAGTGGTC
AATAATGACCACTTTCTCTACTGGGGA–

**Figure S1**

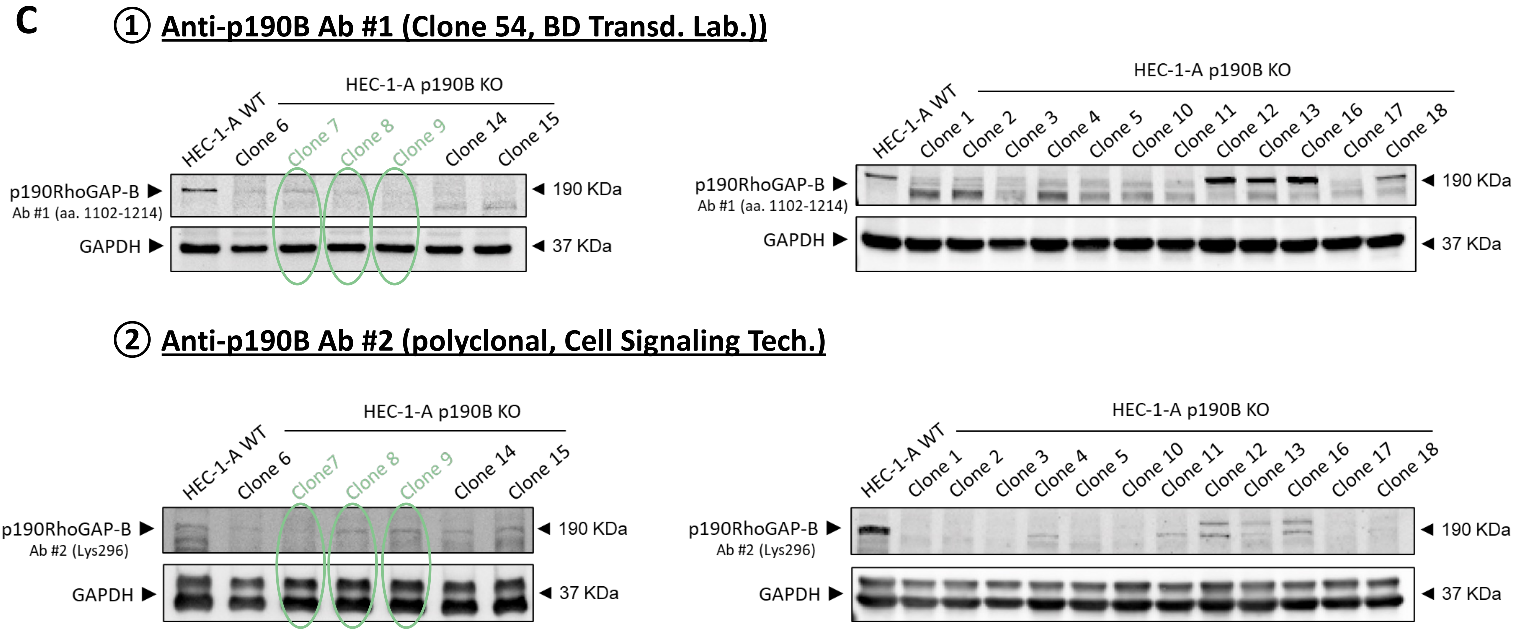


**D**

Clone #7 (KO-B#1)

- **Allele n°1: +1 insertion (green)**

*--**ATG**ATGGCAAAAAACAAAGAGCCTCGTCCCCCATCCCTAT*
*ACCATCAGTATAGTTGGACTCTCTGGGACTGAAAAAGACAAAGGTAACTGTGG*
*AGTTGGAAAGTCTTGTTTGTGCAATAGATTTGTACGCTCA–*

- **Allele n°2: -10 deletion (red)**

*--**ATG**ATGGCAAAAAACAAAGAGCCTCGTCCCCCATC~~CTAT~~*
*~~ACCATC~~AGTATAGTTGGACTCTCTGGGACTGAAAAAGACAAAGGTAACTGTGG*
*AGTTGGAAAGTCTTGTTTGTGCAATAGATTTGTACGCTCA–*

Clone #9 (KO-B#2)

- **Allele n°1: +1 insertion (green)**

*--**ATG**ATGGCAAAAAACAAAGAGCCTCGTCCCCCATCCGTAT*
*ACCATCAGTATAGTTGGACTCTCTGGGACTGAAAAAGACAAAGGTAACTGTGG*
*AGTTGGAAAGTCTTGTTTGTGCAATAGATTTGTACGCTCA–*

- **Allele n°2: -10 deletion (red)**

*--**ATG**ATGGCAAAAAACAAAGAGCCTCGTCCCCCATC~~CTAT~~*
*~~ACCATC~~AGTATAGTTGGACTCTCTGGGACTGAAAAAGACAAAGGTAACTGTGG*
*AGTTGGAAAGTCTTGTTTGTGCAATAGATTTGTACGCTCA–*

Clone #8 (KO-B#3)

- **Allele n°1: -10 deletion (red)**

*--**ATG**ATGGCAAAAAACAAAGAGCCTCGTCCCCCATC~~CTAT~~*
*~~ACCATC~~AGTATAGTTGGACTCTCTGGGACTGAAAAAGACAAAGGTAACTGTGG*
*AGTTGGAAAGTCTTGTTTGTGCAATAGATTTGTACGCTCA–*

- **Allele n°2: -2 deletion (red)**

*--**ATG**ATGGCAAAAAACAAAGAGCCTCGTCCCCCATC~~CT~~AT*
*ACCATCAGTATAGTTGGACTCTCTGGGACTGAAAAAGACAAAGGTAACTGTGG*
*AGTTGGAAAGTCTTGTTTGTGCAATAGATTTGTACGCTCA–*

**Figure S1**

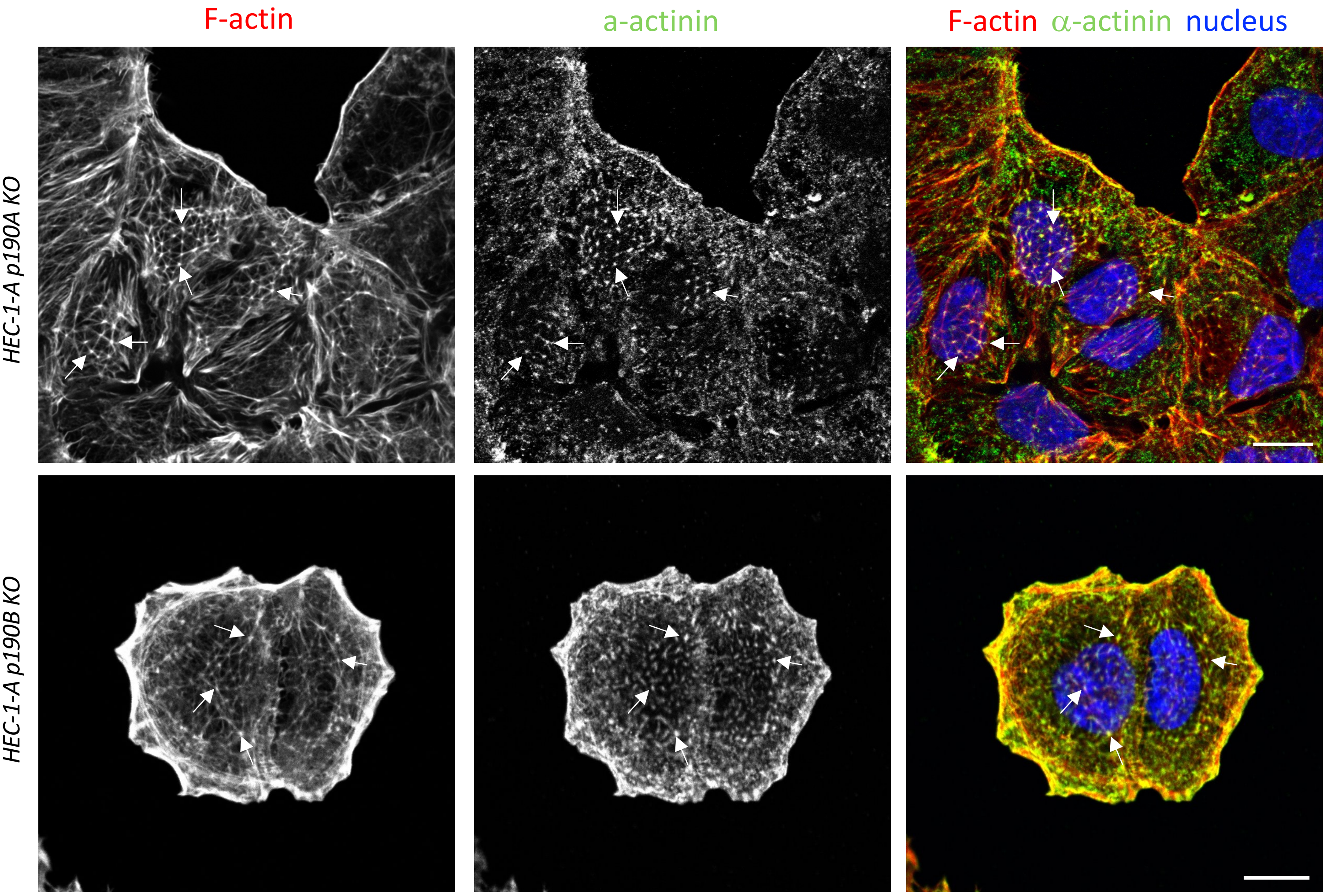
F-actin
a-actinin
F-actin α-actinin nucleus
HEC-1-A p190A KO
HEC-1-A p190B KO

**Figure S2**

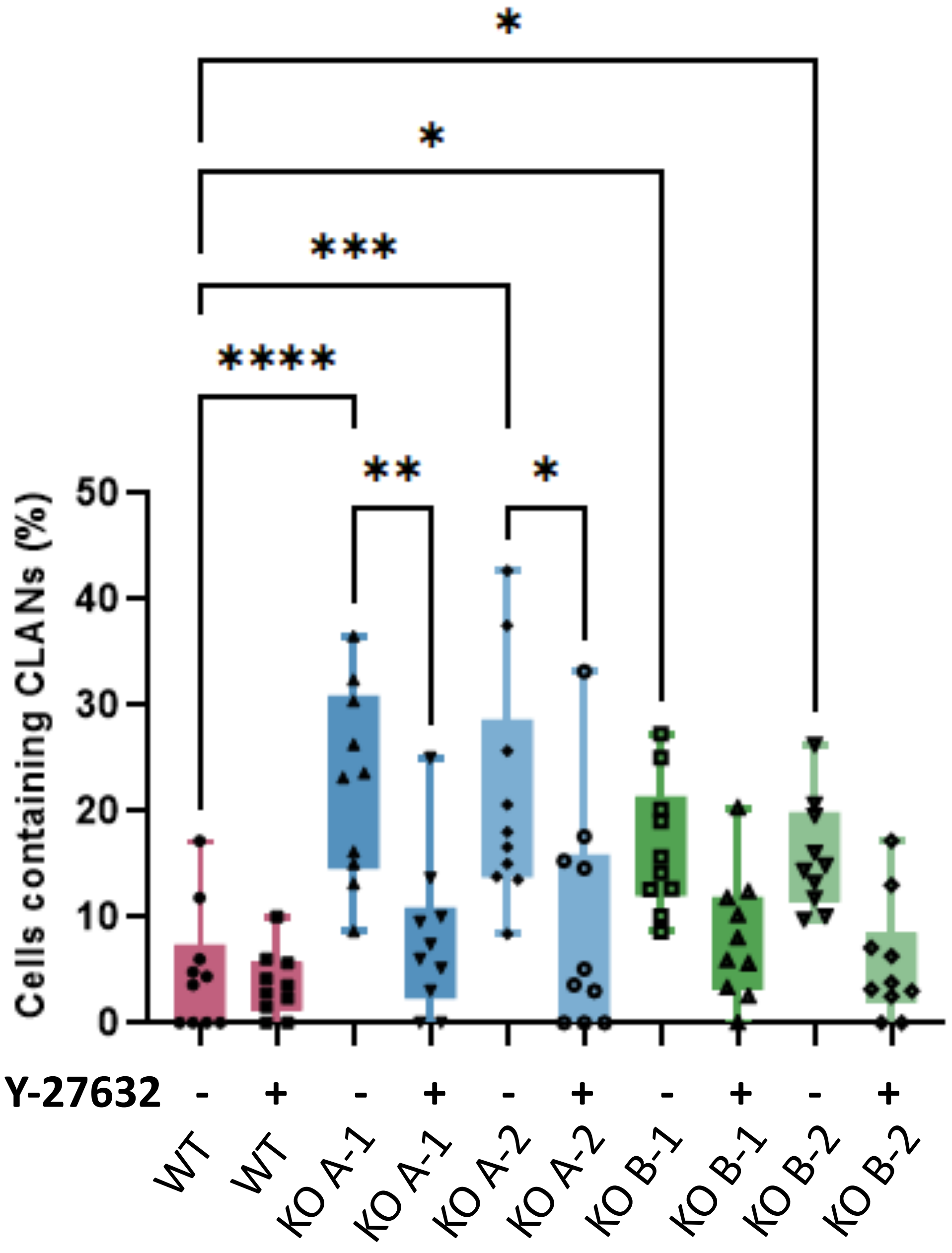
*
*
***
****
**
*
Cells containing CLANs (%)
50
40
30
20
10
0
Y-27632
- + - + - + - + - +
WT
WT
KO A-1
KO A-1
KO A-2
KO A-2
KO B-1
KO B-1
KO B-2
KO B-2

**Figure S3**

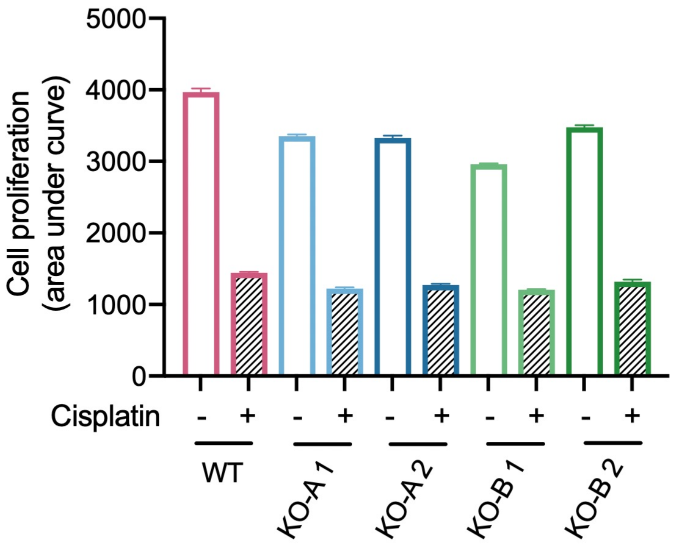
5000
4000
3000
2000
1000
0
Cell proliferation
(area under curve)
Cisplatin
- +
- +
- +
- +
- +
WT
KO-A 1
KO-A 2
KO-B 1
KO-B 2

### Figure S4
