## supplemental tables for "p190A/*ARHGAP35* and p190B/*ARHGAP5* proteins in endometrial cancer, a novel cancer-relevant paralog interplay"

**<u>Supplementary Table 1:</u> Clinical and tumor characteristics of the patients included in this study.** Patients are numbered from P1 to P16. Sample information is indicated as "NT(1)/T(3)," where "T" refers to tumoral tissue, "NT" to non-tumoral tissue, and the number in parentheses denotes the number of samples obtained from each patient. AWD, alive with disease; DOD, died of disease; DWD, died with disease; NED, no evidence of disease.

| Patient | Samples | Cancer type | Stage | Grade | % of normal cells in NT sample | % of cancer cells in T sample | Follow Up | Age |
|---|---|---|---|---|---|---|---|---|
| **P1** | T(1) | Undifferenciated | FIGO stage IA | grade 3 | | 40 | NED | 72 |
| **P2** | NT(1)/T(3) | Endometrioid | FIGO stage II | grade 2 | 60 | 60/70/60 | NED | 44 |
| **P3** | NT(1)/T(1) | Clear cell | FIGO stage IIIC | grade 3 | 20 | 40 | DWD | 78 |
| **P4** | T(3) | Endometrioid | FIGO stage II | grade 2 | | 60/60/50 | NED | 65 |
| **P5** | T(1) | Endometrioid | FIGO stage IA | grade 2 | | 60 | NED | 70 |
| **P6** | NT(1)/T(1) | Serous | FIGO stage IVB | grade 3 | 30 | 70 | DWD | |
| **P7** | T(1) | Endometrioid | FIGO stage IIIC | grade 3 | | 40 | T | |
| **P8** | NT(1)/T(1) | Endometrioid | FIGO stage IB | grade 1 | 30 | 30 | NED | |
| **P9** | NT(1)/T(1) | Endometrioid | FIGO stage IA | grade 1 | 20 | 0 | DOD | |
| **P10** | NT(1)/T(1) | Serous | FIGO stage IB | grade 2 | 60 | 60 | T | |
| **P11** | NT(1)/T(3) | Serous | FIGO stage IA | | 60 | 40/40/60 | NED | |
| **P12** | NT(1)/T(2) | Endometrioid | FIGO stage IB | grade 3 | 50 | 60/60 | T | |
| **P13** | NT(1)/T(1) | Serous | FIGO stage IA | | 40 | 30 | DOD | |
| **P14** | NT(1)/T(1) | Endometrioid | FIGO stage IA | grade 1 | 60 | 50 | NED | |
| **P15** | NT(1)/T(1) | Serous | FIGO stage IV | grade 3 | 60 | 70 | AWD | |
| **P16** | NT(1)/T(3) | Serous | FIGO stage IVB | | 40 | 70/70/70/70 | AWD | |

**<u>Supplementary Table 2</u>: Mutational status of endometrial cell lines for *ARHGAP35* and *ARHGAP5* genes.** Data were extracted from the Cancer Cell Line Encyclopedia (CCLE) database (https://sites.broadinstitute.org/ccle/)

| **Cell lines** | ***ARHGAP35*** | ***ARHGAP5*** |
|---|---|---|
| AN3CA | WT | p.P969H (Missense mutation) |
| HEC-1-A | p.K35E (Missense)<br>p.L351R (missense) | WT |
| HEC-1-B | p.L351R (Missense mutation) | WT |
| Ishikawa | WT | p.F935fs (Frameshift) |
| KLE | WT | p.A1357A (Silent) |
| RL95-2 | WT | p.N155N (Silent)<br>p.F965F (Silent)<br>p.R955S (Missense) |
